# A novel widespread MITE element in the repeat-rich genome of the *Cardinium* endosymbiont of the spider *Oedothorax gibbosus*

**DOI:** 10.1101/2022.06.21.494476

**Authors:** Tamara Halter, Frederik Hendrickx, Matthias Horn, Alejandro Manzano-Marín

## Abstract

Free-living bacteria have evolved multiple times to become host-restricted endosymbionts. The transition from a free-living to a host-restricted lifestyle comes with a number of different genomic changes, including a massive loss of genes. In host-restricted endosymbionts, gene inactivation and genome reduction is facilitated by mobile genetic elements, mainly insertion sequences (ISs). ISs are small autonomous mobile elements, and one of, if not the most, abundant transposable elements in bacteria. Proliferation of ISs is common in some facultative endosymbionts, and is likely driven by the transmission bottlenecks, which increase the level of genetic drift. In the current study we present a manually curated genome annotation for a *Cardinium* endosymbiont of the dwarf spider *Oedothorax gibbosus. Cardinium* species are host-restricted endosymbionts that, similarly to *Wolbachia* spp., include strains capable of manipulating host reproduction. Through the focus on mobile elements, the annotation revealed a rampant spread of ISs, extending earlier observations in other *Cardinium* genomes. We found that a large proportion of IS elements are actually pseudogenised, with many displaying evidence of recent inactivation. Most notably, we describe the lineage-specific emergence and spread of a novel IS-derived **M**iniature **I**nverted repeat **T**ransposable **E**lement (MITE), likely being actively maintained by intact copies of its parental IS982-family element. This work highlights the relevance of manual curation of these repeat-rich endosymbiont genomes for the discovery of novel MITEs, as well as the possible role these understudied elements might play in genome streamlining.

## Introduction

Microbial symbionts are widespread across the animal kingdom, shaping their hosts’ evolution and serving them as a source for novel metabolic capabilities (Douglas, 2014; McFall-Ngai *et al*., 2013). Of particular interest are those relations that evolve between microbes and their hosts where the microbe cannot thrive outside of its host’s cells or tissues. Within these endosymbionts, we find the widely studied obligate nutritional symbioses observed in phloem-or blood-feeders as well as those involving facultative endosymbiotic lineages (Moran *et al*., 2008). A widespread feature of the genomes of facultative endosymbionts is the presence of large numbers of mobile elements; including prophages, group II introns, and mainly insertion sequences (hereafter **IS**s) (Degnan *et al*., 2009, 2010; Latorre and Manzano-Marín, 2017; McCutcheon and Moran, 2012; Patel *et al*., 2019). In their simplest form, ISs are small autonomous transposable elements encoding for a transposase gene flanked by terminal inverted repeats. ISs are arguably the most numerous mobile elements in bacteria (Siguier *et al*., 2014), and have been implicated in promoting widespread genome rearrangement and differential pseudogenisation between closely related strains of facultative endosymbionts (Belda *et al*., 2010; Clayton *et al*., 2012; Gillespie *et al*., 2012; Halter *et al*., 2022a; Manzano-Marín and Latorre, 2014).

Facultative endosymbionts include both conditional beneficial symbionts as well as reproductive manipulators. Reproductive manipulation entails, most famously, male killing, feminization, and cytoplasmic incompatibility (hereafter ***CI***) (Hurst and Frost, 2015) which can facilitate in the spread of the endosymbiont in a host population (Tram *et al*., 2003). While *Wolbachia* strains are the most notorious male-killing and *CI*-inducing endosymbionts, specific strains of the endosymbiotic genus *Cardinium* have also been shown to be involved in inducing *CI* (Hunter *et al*., 2003), parthenogenesis (Zchori-Fein *et al*., 2001), and feminisation (Weeks and Breeeuwer, 2003). Analysis of the genome of a *CI*-inducing *Cardinium* strain from the parasitoid wasp *Encarsia pergandiella* suggests an independent evolution of the CI phenotype in *Wolbachia* and *Cardinium* lineages (Penz *et al*., 2012). Based on available genome sequences, *Cardinium* strains have been organised into three groups, with group **A** containing exclusively endosymbionts from arthropods (namely insects and mites), group **B** from nematodes, and group **C** from *Culicoides punctatus* (Diptera: Ceratopogonidae) (Siozios *et al*., 2019). Similarly to *Wolbachia*, phylogenetic analyses suggest that occasional switching between distant host phyla may be a feature of the genus *Cardinium* (Siozios *et al*., 2019). To date, eight genomes of different finishing status are available in the databases. All strains hold genomes of around 1 Mega base-pair (**Mbp**) and are highly enriched in mobile elements, namely insertion sequences (hereafter **ISs**). Despite the important role IS elements play in both genome inactivation and genome rearrangement, only the cBtQ1 strain isolated from the whitefly *Bemisia tabaci* biotype MEDQ1 has undergone rigorous annotation of these elements (Santos-Garcia *et al*., 2014). In the present work, we present the manually curated annotation of the *Cardinium* endosymbiont of the spider *Oedothorax gibbosus*. This revealed an abundant small non-autonomous IS-derived Miniature Inverted repeat Transposable Element (or MITE), which to our knowledge, is previously unreported for endosymbionts and is unique to this *Cardinium* strain.

## Results and Discussion

The genome of *Cardinium* strain cOegibbosus-W744×776 (hereafter cOegib) was assembled previously from long- and short-read data generated for the genome sequencing of its spider host, *Oedothorax gibbosus* (Halter *et al*., 2022b; Hendrickx *et al*., 2021). To produce a high-quality annotation of the mobile elements of cOegib, an initial draft annotation was done using Prokka v1.14.6 (Seemann, 2014). This draft annotation was followed by careful manual curation using a combination of DELTA-BLASTP (*vs*. NCBI’s nr and Swiss-Prot) (Boratyn *et al*., 2012), InterProScan v5.45-80.0 (Jones *et al*., 2014), Infernal v1.1.3 (–cut tc –mid; vs. Rfam v14.2) (Kalvari *et al*., 2021; Nawrocki and Eddy, 2013), tRNAscan-SE v2.0.9 (-B –isospecific) (Chan *et al*., 2021), and ARAGORN v1.2.38 (Laslett, 2004). Finally, careful manual searches against the ISfinder database (Siguier *et al*., 2006) were performed in order to identify complete, partial, and fragmented elements across the genome, with special care to correctly identify the terminal inverted repeats of IS elements.

As previously reported in Halter *et al*. (2022b), the general features of the genome of cOegib are comparable to other members of the *Cardinium* genus (supplementary table S1). Similarly to other *Cardinium* strains (Penz *et al*., 2012; Santos-Garcia *et al*., 2014), a large fraction of cOegib’s genome (29.60%) is made up of mobile elements, which is the highest reported for the genus. Manual curation revealed that group II introns and ISs made up the majority of these elements (87.97%), with the latter being by far the most abundant. From the repertoire of in total 300 ISs, we were able to identify ISCca2 to ISCca6 elements, previously reported for other *Cardinium* strains (Penz *et al*., 2012; Santos-Garcia *et al*., 2014). Nonetheless, their copy numbers are dissimilar to those of strain cBtQ1, revealing lineage-specific expansions/contractions (table 1). In addition to these *Cardinium*-specific ISs, we were able to detect an additional eight mobile elements (named ISCca9 to ISCca16) belonging to seven different IS families. Despite being highly abundant, only 23.33% of these IS elements, belonging to six different types, encode for an intact transposase gene, and 8.00% are only partial IS elements with no transposase gene/pseudogene. This suggests that only the subset of IS elements that have at least one intact copy in the genome still preserve the ability to mobilise, while those that do not are likely to be eventually purged, unless new intact copies are acquired through horizontal gene transfer. This hypothesis is supported by the five most abundant IS elements matching all but one of the intact ones.

**Table 1.**
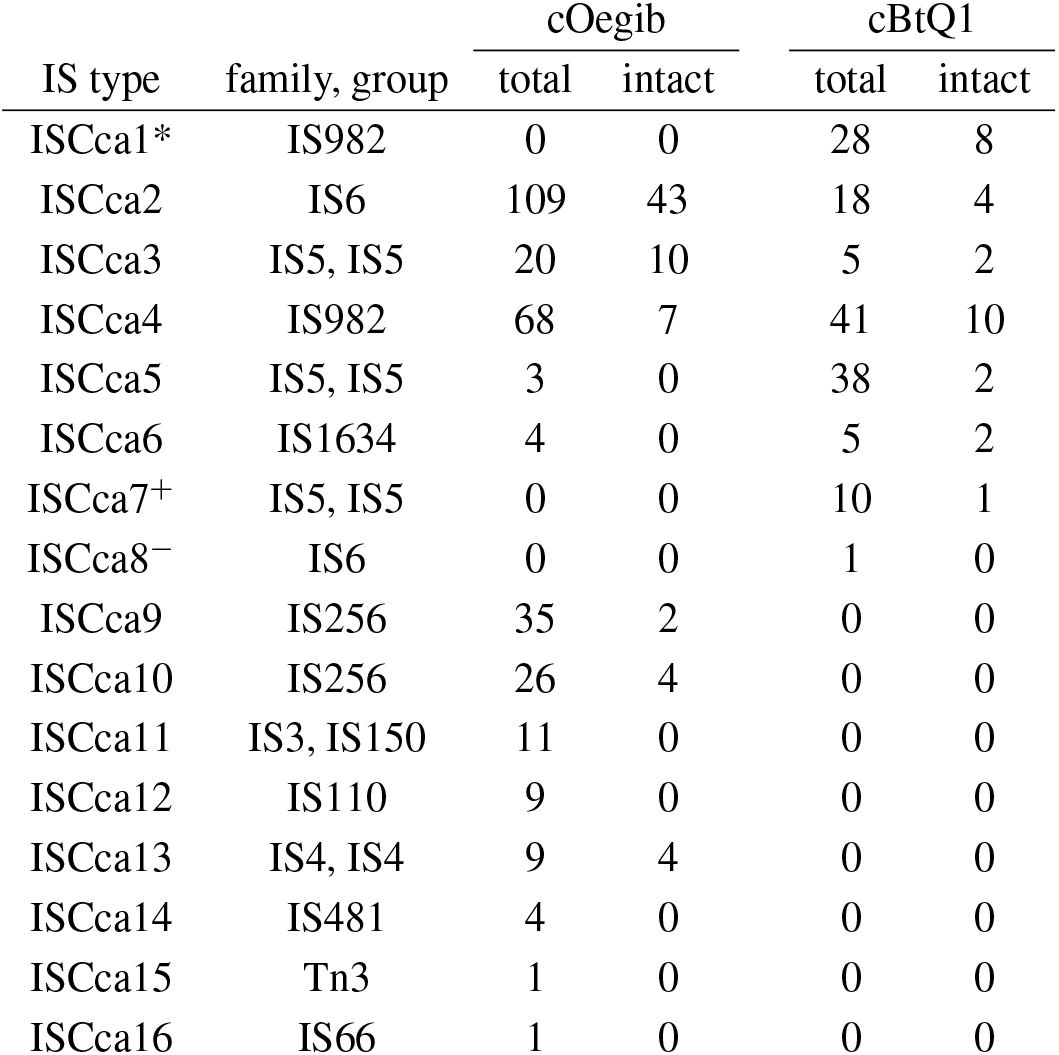
Distribution of intact and partial IS elements in selected Cardinium strains. * ISCca1 is highly similar to ISCca4. ^+^ ISCca7 is highly similar to ISCca3. ^−^ ISCca8 is highly similar to ISCca2. These two Cardinium strains were selected given that they are the only two to undergo thorough manual curation for IS-elements.

Most notably, manual curation of the IS elements revealed a large number of a shorter sequence of *circa* 240 base pairs (**bp**) flanked by very similar inverted repeats to those of ISCca4 (IS982 family; figure 1A), the second most abundant IS element in the cOegib genome. However, these shorter repeats completely lack a transposase gene. Upon closer inspection, we found good evidence to suggest that these shorter repeats are likely derived from a parental ISCca4 element: downstream of the left inverted repeat,there is a short 5 bp-long conserved sequence when compared to ISCca4. This novel *Cardinium* IS-like element, designated MITECca01, has all the features of **M**iniature **I**nverted repeat **T**ransposable **E**lements (or **MITE**s), which are short (typically shorter than 300 bp) non-autonomous transposable elements that depend on a functional transposase gene of their parental IS to mobilise (reviewed in Siguier *et al*., 2014). Hitherto, MITEs have been reported in few plants, bacteria, and archaea, with no reports, to our knowledge, in maternally-inherited intracellular endosymbionts. This under-reporting might be due to their small nature, a lack of thorough annotation of mobile genetic elements in genomes, and the lack of available full-length annotations (i.e. including terminal inverted repeats) of their parental IS elements.

**Figure 1.**
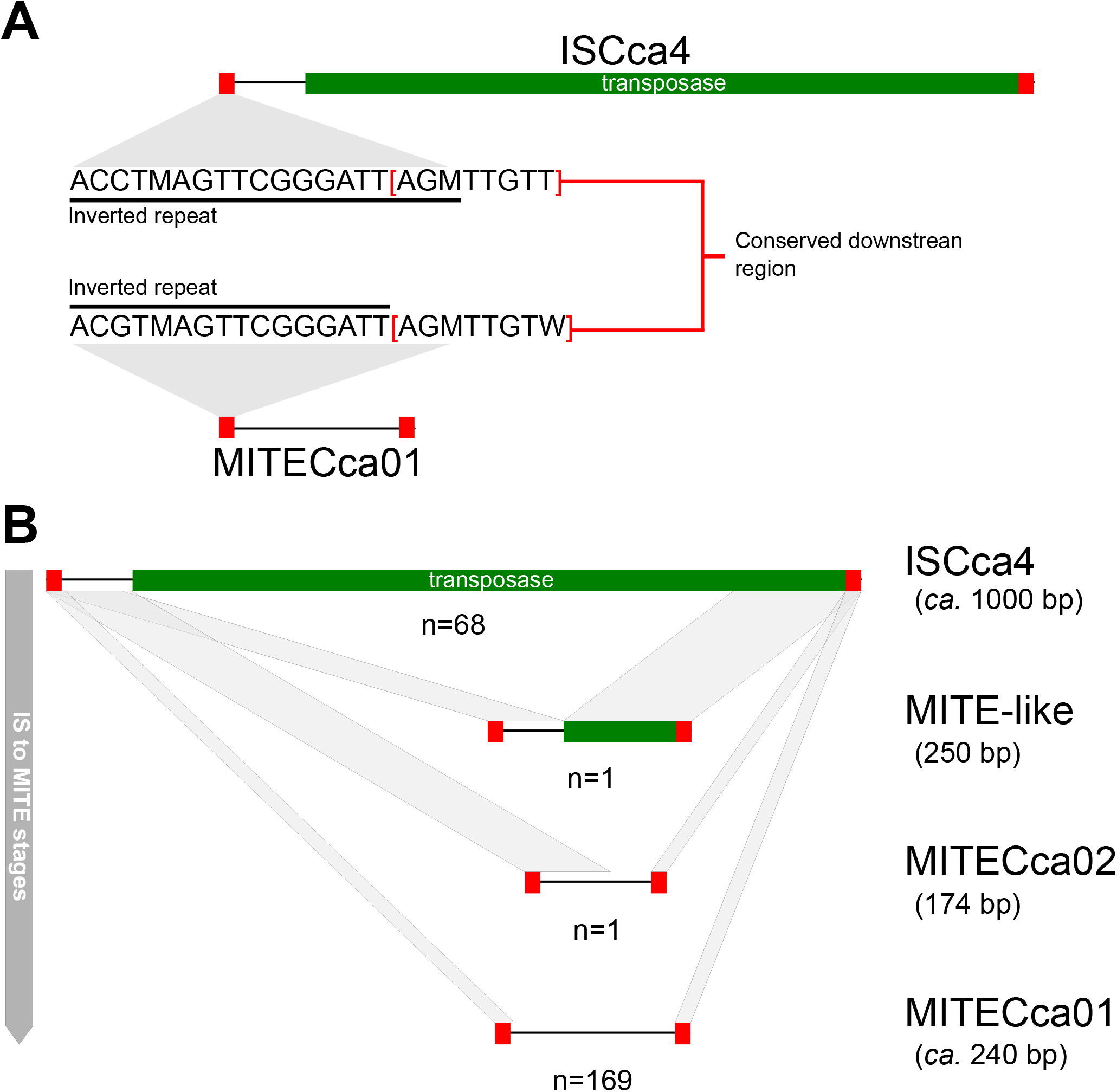
Diagram depicting ISCca4, MITE-like, and MITE elements detected in cOegib. **(A)** Comparison of ISCca4 and MITECca01 and their inverted repeats. **(B)** Diagram depicting, form top to bottom, the ISCca4 and related MITE elements in order of likely stages of MITE-element formation and copy number increase across the genome. At the bottom-right of each panel, the number of copies is expressed as “n” under the diagram of each element.

The newly identified MITE has successfully spread throughout the genome of cOegib, effectively becoming the most abundant mobile element in the genome, with 169 copies and making up 3.5% of the total chromosomal sequence. This large number of copies contrasts even the five most abundant IS elements, which are present in 109 (ISCca2), 68 (ISCca4), 35 (ISCca9), 26 (ISCca10), and 20 (ISCca3) copies, respectively. Similarly to other ISs in the genome of cOegib and other endosymbionts (Belda *et al*., 2010; Clayton *et al*., 2012; Manzano-Marín and Latorre, 2014) MITECca01 was found mostly in intergenic regions as well as disrupting other mobile elements, and less commonly inactivating protein-coding genes. Its genomic distribution strongly points towards a role for MITECca01 in IS element inactivation in the genome of cOegib. Contrary to IS elements, very few MITECca01 elements are found disrupted by other mobile elements, which could likely be due to their small size compared to ISs (240 *vs. ca*. 1000 bp). Further, we were able to identify a second MITE element very similar to MITECca01, termed MITECca02 (figure 1B), which likely represents yet another MITE element with potential to spread throughout this *Cardinium* lineage. This second MITE element might be evolutionarily younger, given its copy number and the fact it keeps a much longer conserved sequenced with ISCca4 following the left inverted repeat (supplementary file S1). On top of these two MITE elements, we also identified an uninterrupted highly eroded ISCca4 element of 250 bp in length that retains a small part of the 3’-end of its transposase gene (supplementary file S1). This IS remnant potentially represents a very early stage of the birth of a MITE element. Finally, blastn searches of these novel *Cardinium* MITE elements did not reveal the presence of these in any of the other sequenced *Cardinium* strains, suggesting that it originated in the lineage leading to cOegib.

In conclusion, through the thorough annotation of the *Cardinium* strain cOegib we were able to shade light on the dynamics that likely resulted in the current distribution and abundance of mobile elements in this genome. In addition, the MITECca01 element present in the cOegib genome represents, to our knowledge, a novel kind of IS-like element in a maternally-inherited endosymbiont lineage. This novel MITE has been very successful in multiplying across the genome, and its mobility and persistence is likely linked to the survival of intact copies of the “parental” ISCca4 element. Finally, the genome of cOegib along with its careful and detailed annotation represents a valuable resource with relevance to the continued study of this endosymbiont taxon and transposable elements, as well as the larger field of reductive genome evolution.

## Supplementary Material & Data Availability

Supplementary table S1 and file S1 have been included in this submission. The genome annotation of the *Cardinium* endosymbiont of *Oedothorax gibbosus* strain cOegibbosus-W744×776 has been deposited in the European Nucleotide Archive (ENA) under accession number OW441264.

## Acknowledgements

This project has received funding from the University of Vienna (uni:docs to T.H.) and the *Austrian Science Fund FWF* (DOC 69-B). A.M.M. was supported by the European Union’s Horizon 2020 research and innovation programme under a *Marie Skłodowska-Curie Individual Fellowship* (LEECHSYMBIO, grant agreement no. 840270). The funders had no role in study design, data collection and analysis, decision to publish, or preparation of the manuscript.

## Notes

### Competing Interest Statement

The authors have declared no competing interest.

